# Altered heterosynaptic plasticity impairs visual discrimination learning in adenosine A1 receptor knockout mice

**DOI:** 10.1101/2020.11.17.386912

**Authors:** Renee Chasse, Alexey Malyshev, R. Holly Fitch, Maxim Volgushev

## Abstract

Theoretical and modeling studies demonstrate that heterosynaptic plasticity - changes at synapses inactive during induction - facilitates fine-grained discriminative learning in Hebbian-type systems, and helps to achieve a robust ability for repetitive learning. A dearth of tools for selective manipulation has hindered experimental analysis of the proposed role of heterosynaptic plasticity in behavior. Here we circumvent this obstacle by testing specific predictions about changes in heterosynaptic plasticity, and associated behavioral consequences, following experimental manipulation of adenosine A1 receptors (A1R). We show that, compared to wild-type controls, A1R-knockout mice have impaired synaptic plasticity in visual cortex neurons, coupled with significant deficits in visual discrimination learning. Deficits in A1R-knockouts were seen specifically during re-learning, becoming progressively more apparent with learning on sequential visual discrimination tasks of increasing complexity. These behavioral results confirm our model predictions, and provide the first experimental evidence for a proposed role of heterosynaptic plasticity in learning.

**Highlights:**

- Synaptic plasticity is impaired in visual cortex neurons in adenosine A1R knockout mice
- Homosynaptic and heterosynaptic plasticity in A1R KO mice is dominated by depression
- Learning on sequential, increasingly complex visual tasks is impaired in A1R KO mice
- Learning deficits match predicted effects of impaired heterosynaptic plasticity

## INTRODUCTION

Adenosine is an abundant activity-dependent metabolite of ATP and a potent endogenous neuromodulator. Adenosine is involved in regulation of sleep homeostasis and slow-wave sleep oscillations, mediation of negative feedback in response to excessive activity, and neuroprotection from ischemia or hypoxia (Mendonҫa et al., 2000; Dunwiddie and Masino 2001; Bjorness and Greene 2009; Halassa et al 2009; Cunha 2005.

In cortical neurons, activation of adenosine A1 receptors (A1Rs) suppresses synaptic transmission, and modulates long-term plasticity in hippocampus (Mendonҫa et al., 2000; Moore et al., 2003; Izumi and Zorumski 2008; Dias et al., 2013;. Pérez-Rodríguez et al., 2019) and neocortex (Blundon et al., 2011; Bannon et al., 2017). In layer 2/3 pyramidal neurons from rat visual cortex, blockade of A1Rs led to a decrease in the proportion of inputs expressing LTP, and an increase of the proportion of inputs expressing LTD (Bannon et al., 2017). This shift toward depression was observed for *both* synapses activated during induction (homosynaptic plasticity), as well as synapses not activated during induction (heterosynaptic plasticity). In model neurons, experimentally-observed A1R-modulation of heterosynaptic plasticity could shift their operating point along a continuum, from a regime of predominantly associative plasticity to predominantly homeostatic regime (Bannon et al., 2017). In the homeostatic regime, synapses with excessively increased or decreased weights are brought back into operational range, and the system is prepared for subsequent learning. Blockade of A1Rs disrupted this homeostatic regime, and impaired new learning in model neurons (Volgushev et. al., 2016). Modelling results also predict that impairment of homosynaptic versus heterosynaptic plasticity should lead to different learning deficits. Learning deficits caused by impairment of homosynaptic associative plasticity should be evident already during initial stages of learning, but could be mild unless associative plasticity is completely blocked or impaired severely. In contrast, impairment of heterosynaptic plasticity may not impair initial learning, but would specifically disrupt subsequent learning and re-learning (e.g., task reversals). Such learning deficits should become progressively more apparent with successive learning tasks. Here, we tested these differential predictions using A1R −/− knockout (A1R-KO) mice.

Prior research shows that in the hippocampus, synaptic plasticity is impaired during acute blockade of A1Rs (Mendonҫa et al., 2000; Moore et al., 2003; Izumi and Zorumski 2008; Dias et al., 2013;. Pérez-Rodríguez et al., 2019), yet no difference is seen between A1R-KO and wild type (WT) animals in synaptic plasticity or spatial learning (Giménez-Llort et. al., 2002; 2005). Therefore, we first asked whether synaptic plasticity in visual cortex is different in A1R-KO and WT animals. Second, we tested specific predictions about differential learning deficits in A1R-KO and WT mice using a series of progressively more difficult visual discrimination tasks.

## RESULTS

We used A1R-KO (−/−) and littermate WT mice from a breeding colony at the University of Connecticut (B6N.129P2-*Adora1*^tm1Bbf^/J mice, *The Jackson Laboratory*).

### Impaired synaptic plasticity in visual cortex neurons of A1R knockout mice

We first tested for differences in synaptic plasticity in visual cortex neurons from A1R-KO and WT mice. In layer 2/3 pyramidal neurons, we recorded small amplitude EPSPs evoked by electrical stimulation in layer 4. In slices from WT animals, pairing synaptic stimulation with bursts of postsynaptic spikes (Fig. 1 inset) typically induced long-term potentiation. In a sample neuron (Fig. 1a), EPSP amplitude increased after pairing to 154% of control. This pairing procedure induced LTP in 5 WT neurons, LTD in 3, and in one cell no changes were observed. On average, EPSP amplitude after pairing was 113.7±12.4% of control (Fig. 1c, N=9). By contrast, in slices from A1R-KO animals, the same pairing procedure induced long-term depression. In a sample neuron (Fig. 1b), EPSP amplitude decreased after pairing to 57.2% of control. LTD was observed in 9 neurons, LTP in only 3 cases, and in the remaining 5 cells EPSP did not change. On average, EPSP amplitude decreased after pairing in A1R-KO animals (Fig. 1f, 84.6±6.0% of control, N=17, p=0.051). The differential effects of pairing on synaptic transmission in WT and KO animals were reflected in differences in the average EPSP amplitude changes (Fig. 1d, 113.7±12.4% in WT *vs.* 84.6±6.0% in KO, p=0.033), and in the higher frequency of LTP in WT *versus* LTD in KO neurons (Fig. 1e).

**Figure 1.**
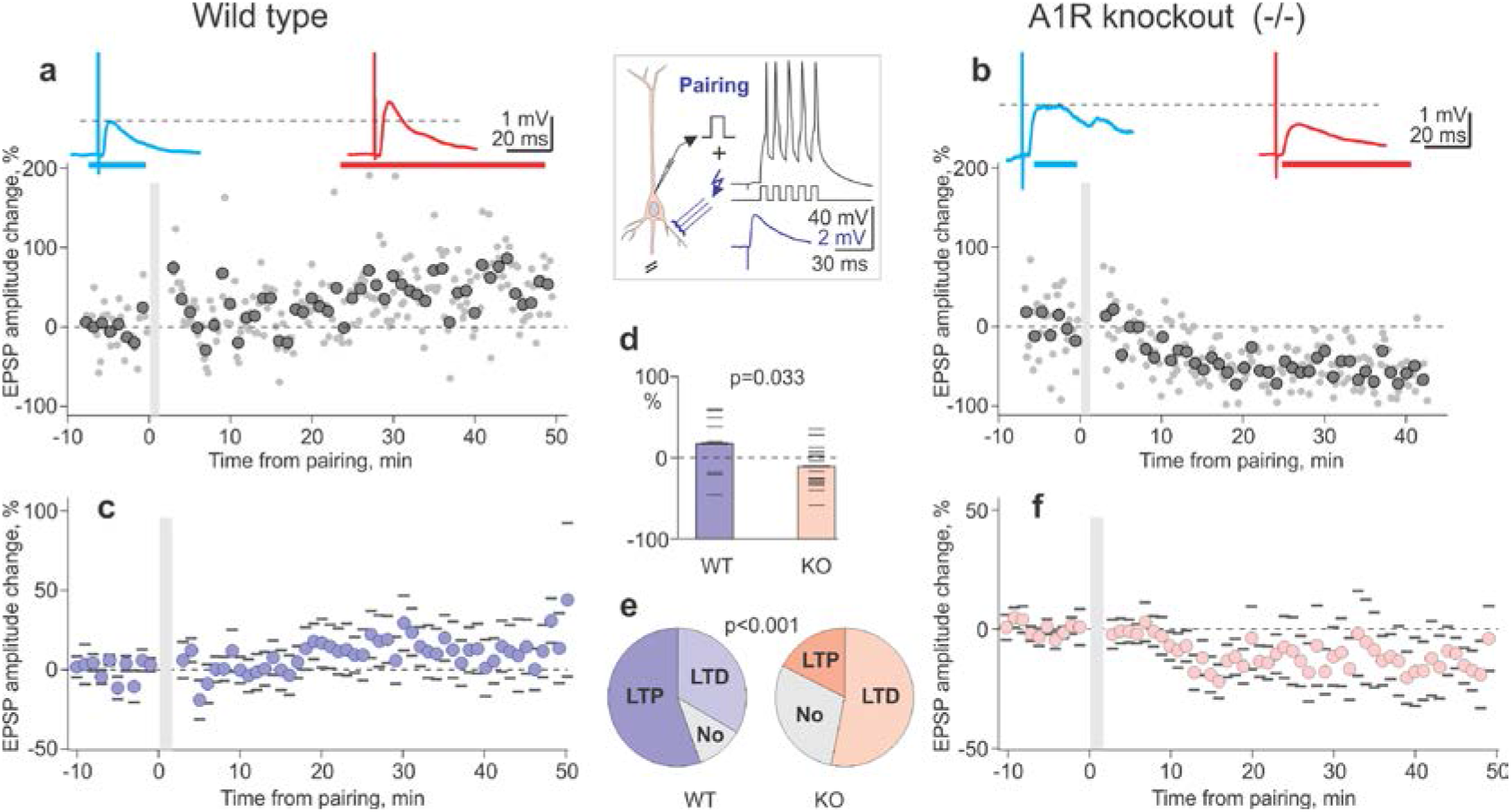
Impaired long-term potentiation in L2/3 neurons from visual cortex of A1R-KO mice. **a, b**: Pairing procedure (inset, synaptic stimulation followed with a 10 ms delay by 5 action potentials at 100 Hz, repeated 30 times) typically induced long-term potentiation (LTP) in neurons from WT animals (**a**) but long-term depression (LTD) in neurons from A1R-KO mice (**b**). Time course shows individual EPSP amplitudes (dots) and averages over 1 minute (large symbols), before and after the pairing procedure (grey vertical bar). EPSPs above the time course are averages over the indicated periods. **c, f**: Summary time course of EPSP amplitude changes in N=9 neurons from wild type (**c**) and N=17 neurons from A1R-KO mice (**f**). Averages over 1 minute with SEM. **d**: Pairing-induced changes of EPSP amplitudes in individual neurons (horizontal dash symbols) and averaged data (bars) for WT and A1R-KO mice (T-test, p=0.033). **e**: Frequency of occurrence of LTP and LTD after pairing procedure in neurons from WT and A1R-KO mice (Chi-Square test, p<0.001).

Notably, heterosynaptic plasticity induced by intracellular tetanization (Fig. 2a; Bannon et. al., 2017; Volgushev et. al., 2000; 2016) was impaired in KO animals in the same way as pairing-induced homosynaptic plasticity. In WT animals, intracellular tetanization induced LTP in 8, LTD in 7, and led to no changes in 7 inputs. Heterosynaptic changes were balanced, with averaged amplitude of 102±6.8% (N=22) of control (Fig. 2b, 2c). In contrast, heterosynaptic depression dominated in KO animals, with LTD in 13, LTP in only one, and no changes in 6 inputs (difference from WT: p<0.001). On average, the EPSP amplitude in KO animals was depressed by intracellular tetanization to 74.5+5.0% of control (N=20, p=0.019; difference from WT: p=0.0027).

**Figure 2.**
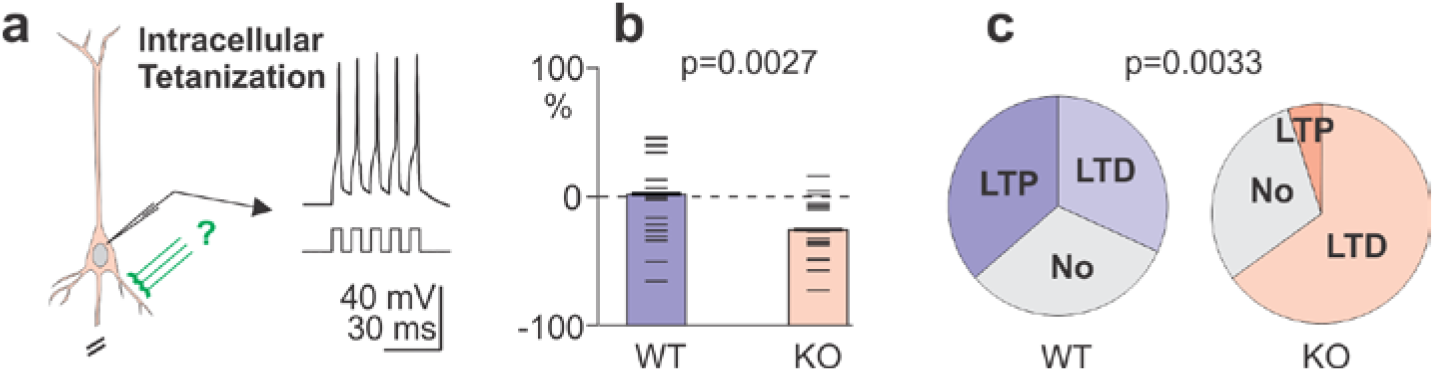
Impaired heterosynaptic plasticity in L2/3 neurons from visual cortex of A1R-KO mice. **a**: A scheme of intracellular tetanization, bursts of depolarization-induced postsynaptic spikes without presynaptic activation. **b**: EPSP amplitude changes after intracellular tetanization in WT (N=22 inputs) and KO (N=20) animals. **c**: Frequency of occurrence of LTP and LTD after intracellular tetanization in neurons from WT and KO mice.

Overall, induction of both homosynaptic and heterosynaptic plasticity in KO animals was impaired. This disruption in KO animals manifested as a shift of the balance between LTP and LTD, toward a higher proportion of LTD relative to WT animals.

### Impaired visual discrimination learning in A1R knockout mice

Next, we tested whether altered plasticity in visual cortex neurons of KO mice was associated with changes in the ability to learn progressively more difficult visual tasks. Behavioral operant testing was performed on N=18 A1R-KO (13 female, 5 male) and N=30 WT (11 female, 19 male) animals, blind to genotype, using automated Bussey-Saksida touchscreen chambers (*Lafayette Instruments*, Lafayette, IN). Large (about 10% of the screen area), high contrast (75%) clearly distinct geometric shapes were presented in the lower part of a touchscreen (30.7 cm, resolution 800×600), within reach of the subject. Touching the stimulus on the screen activated food delivery (Strawberry Ensure Plus, *Abbott Nutrition,* Columbus, OH) in the reward tray located opposite the screen. During the period of behavioral testing, subjects were food-restricted to 85% of baseline weights. Throughout operant training, subjects had one test session per day (Monday-Friday), which lasted either 60 min or until 30 correct responses were rewarded with food.

During first two weeks (pre-training), all mice learned to associate screen presses with reward delivery. After pre-training, subjects learned three visual tasks of increasing difficulty, with progressively higher cognitive demand in the association between visual stimuli and food reward.

In stage 1 task (‘must initiate’, 5 days) subjects learned to initiate presentation of a visual stimulus on the screen by nose-poking, exiting the reward tray, and touching the stimulus to obtain food reward. At this stage, touching non-stimulus parts of the screen had no effect. Both WT and KO mice quickly learned this task. All subjects completed the maximum (N=30) rewarded trials during the first session, and on days 2-5 continued to max-out rewarded trials with few exceptions. There were no differences in the number of rewards obtained by WT and KO animals on any single day (e.g., 29.3±0.47 *vs.* 30.0±0.0; p=0.24 on day 5), nor overall (29.4±0.38 *vs.* 29.1±0.24; p=0.53). Total time to complete the 30 trials was comparable for WTs and KOs (e.g., 1537±148s *vs.* 1510±156s, p=0.91 on day 5).

In stage 2 task (‘punish incorrect’, 5 days) subjects learned to touch only the stimulus and no other part of the screen. Touching the stimulus (‘correct’) was rewarded with food; touching any other portion of the screen (‘incorrect’) was punished by a time-out of 5 seconds (no inputs registered and chamber brightly illuminated, ~60 lx). Both WT and KO subjects rapidly learned this second task. In both groups, the number of correct responses increased from day 1 to 2 (27.4±0.49 to 30.0±0.03, p<0.001 in WT; and from 25.2±0.99 to 28.8±1.02, p<0.001 in KO) and plateaued over days 3-5 (Fig. 3d and 3f, “Stage 2”). For both groups, percent correct responses were near-ceiling on day 1 (91.2±1.6% in WT and 84.8±3.4% in KO), and remained high on days 2-5 (>91.5% for WT and >88% for KO; Fig. 3a and 3c, “Stage 2”). While A1R-KO subjects performed with a very high rate of correct responses, WT subjects were slightly better. Pooled over 5 days, WT subjects made more correct (29.3±0.15 WT *vs.* 27.9±0.43 KO, p<0.001), fewer incorrect (2.57±0.21 WT *vs.* 3.99±0.37 KO, p<0.001) and a higher percent of correct responses (92.3±0.6% WT *vs.* 87.8±1.1% KO, p<0.001). They also completed the training sessions faster (1330±49s WT *vs.* 1825±101s KO, p<0.001).

**Figure 3.**
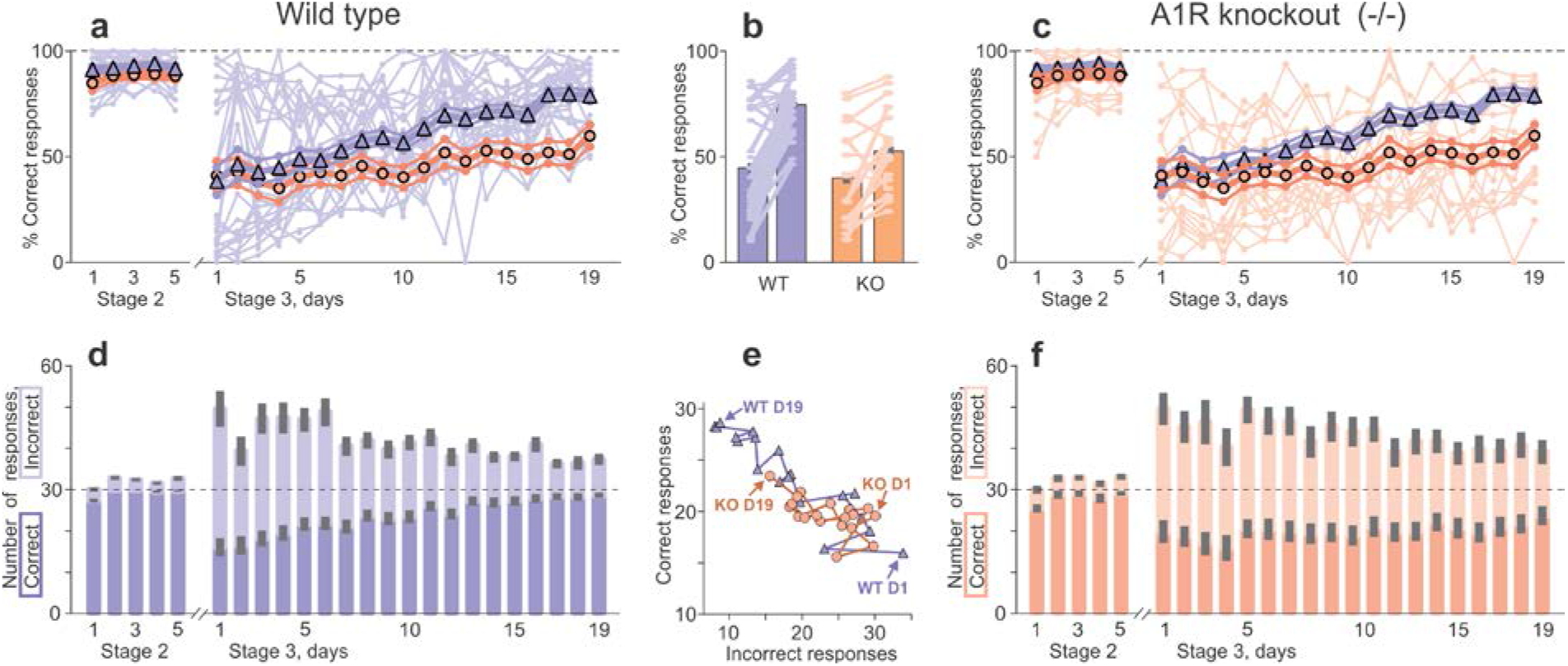
Learning on a difficult but not on a simple visual task is impaired in A1R-KO mice. **a, c**: Percent of correct responses on consecutive testing days 1-5 of learning to touch one stimulus presented on the screen for food reward (stage 2) and days 1-19 of pairwise discrimination, learning to touch correct stimulus out of the two presented (stage 3). Pale lines and dot symbols show data for each WT (**a**) and KO (**c**) subject. Large symbols and thick lines show daily averages with SEM for N=30 WT (triangles, blue) and N=18 A1R-KO (circles, orange) animals. **b**: Percent of correct responses during days 1-6 and days 14-19 of pairwise discrimination learning; Averages for WT (blue) and KO (orange) groups, and data for each subject (connected pale symbols). **d, f**: Number of correct (darker color) and incorrect (lighter color) responses with SEM (grey bars) on consecutive testing days of stags 2 and 3 learning in WT (**d**) and KO (**f**) groups. Horizontal dashed line shows maximal possible number (30) of correct responses per one day. **e**: Number of correct responses plotted against number of incorrect responses on each day of pairwise discrimination task (stage 3, days 1-19) for WT (blue triangles and line) and KO (orange circles and line) groups. Arrows indicate data from the first (D1) and the last (D19) day of testing for WT and KO groups; Lines connect data points from consecutive days.

This difference between WT and KO mice became more pronounced in stage 3 (‘visual pairwise discrimination’, 19 days). In this phase, subjects initiated a trial in which two visually distinct stimuli were presented on screen (randomized left/right position, balanced target assignment). Only touching the correct stimulus was rewarded with food. Touching the incorrect stimulus or blank part of the screen was punished (5 seconds, no inputs registered and bright light ~60 lx in the chamber).

On the first day of the new task, correct responses decreased dramatically (relative to stage 2), and incorrect responses increased (Fig. 3d, 3f; correct: 16.0±2.2 in WT and 19.6±2.8 in KO; incorrect: 33.8±4.4 in WT and 30.0±4.0 in KO). Consequently, percent correct responses decreased (38.4±6.5 % in WT and 41.1±6.8 % in KO; Fig. 3a, 3c). Total time to complete the session dramatically increased compared to stage 2 (2765±209s in WT and 2800±239s in KO animals). However, values did not differ for WT vs. KO on day 1 (p>0.3 for any comparison), indicating that all mice learned from the same baseline.

During subsequent days, WT mice showed clear and consistent learning. The number of correct responses increased over days, and from day 4+ were significantly higher than day 1. Incorrect responses decreased significantly compared to day 1, as seen on days 2, 4, 5 and 7-19 (Fig. 3d, “Stage 3”). Percent correct responses increased, and from day 7+ were significantly higher than on day 1 (Fig. 3a, “Stage 3”). Time to complete the session significantly decreased by day 11, and reached 1695±162s on the last day of training. All of these measures indicate robust learning.

In contrast to WT subjects, KO mice learned much more slowly and less consistently (Fig. 3c; 3f). Out of the three response parameters (correct, incorrect, and percent correct responses), learning was most evident by a decrease in incorrect responses. Compared to day 1, incorrect responses were significantly lower on days 12,14,15 and 17-19 (near-significant on days 13 and 16; p=0.061 and p=0.055). Correct responses tended to increase, but were not significantly higher than day 1 for any of the test days (2-19; Fig. 3f, “Stage 3”). However, mean correct responses during the *last* six days of training were higher than during the *first* six days (days 14-19: 21.2±2.2 vs. days 1-6: 18.4±2.4, p=0.017). Percent correct responses also increased, from 41.1±6.8% on day 1 to 59.9±5.2% on day 19, (Fig. 3c, p=0.012), and from 40.3±5.5 % on days 1-6, to 52.7±4.7% on days 14-19 (Fig. 3b, p<0.001).

Better learning in WT than in KO subjects was clear already during training, but became very pronounced in the last phase of testing. Daily comparisons revealed that, compared to the KO group, WT subjects had significantly higher number of correct (days 11, 13-19; lower number of incorrect (days 9, 13-15, 17-19) and a higher percentage of correct responses (days 8-19; Fig. 3). Over the last six days (14-19) of testing, WT subjects were better than KO on correct (27.9±0.9 *vs.* 21.2±2.2, p=0.0019), incorrect (10.9±1.3 *vs.* 19.7±2.7, p=0.002), and percent correct responses (75.1±2.4% *vs.* 52.7±4.7%, p<0.001; Fig. 3b). On days 14-19 WT animals were also faster to complete sessions (1925±154s *vs*. 2759±225s for KOs (p=0.0029). These results point to a robust impairment of learning in A1R-KO mice as compared to WT animals on the pairwise discrimination task.

Linear model analysis confirmed that Genotype was the main predictor of the observed difference in learning. A linear model considered Percent Correct responses during the final days 14-19 as a response variable. Predictor variables included (i) Genotype, (ii) Sex, (iii) Age, and 5 factors (iv-viii) reflecting performance on stage 2 and the first 6 days of stage 3: (iv) percent correct responses on last day of task 2; (v) number of correct, (vi) number of incorrect, (vii) percent correct and (viii) total number of responses on days 1-6 of task 3. Combinations of predictors optimized to minimize residual standard error *always* included Genotype (function *regsubsets*, R version 3.4.0 (2017-04-21) *The R Foundation for Statistical Computing*). In the linear model that included all predictors (F_DF(8,39)_= 4.856, p<0.001), the only significant predictor of final performance on task 3 was Genotype (t = 3.831, p<0.001 for Genotype; p>0.1 for all others).

In summary, testing on visual tasks of increasing difficulty revealed that both WT and A1R-KO mice could learn the first, most simple task, equally well. Both groups also learned well on the second, more difficult task, though WT animals started to outperform KO subjects. Impairment of learning in KO subjects became clear and pronounced on the third, most difficult task of pairwise discrimination. These results confirm our hypotheses – both the general hypothesis that learning in A1R-KO mice is impaired compared to WT animals, as well as the specific hypothesis that impairment of visual learning in A1R-KO mice becomes progressively more pronounced with increasing task demand.

Interestingly, while learning on the pairwise discrimination task was impaired in KO animals, learning strategies appeared similar in both KO and WT groups. Incorrect responses decreased and correct responses increased for both groups during learning (Fig. 3d and 3f), largely in parallel (Fig. 3e). Moreover, in both groups the reduction in incorrect responses was more pronounced than the increase in correct responses, contributing heavily to increases in percent correct responses. Despite this similarity of strategies, KO animals learned slower and lagged behind WT subjects by several days.

### Baseline motor function, anxiety and locomotor activity are not different in KO and WT mice

Prior studies have reported decreased muscle strength and increased anxiety in A1R-KO mice compared to WT controls (Giménez-Llort et. al., 2002). While several lines of evidence indicate that observed KO impairments (above) were highly task-specific (see Discussion), we nonetheless tested subjects on additional tasks to exclude possible confounds. Assessment for motor function, anxiety and locomotion using a rotarod, elevated plus maze and open field did not reveal any differences between WT and KO animals. On the rotarod test, latency to fall was equivalent in both groups on day 1, and increased on the day 2 (Fig. 4a), indicating comparable motor function and motor learning. On the elevated plus maze test, WT and KO animals spent the same proportion of time on the open arm, indicating no differences in anxiety (Fig. 4b). Results of the open field test likewise showed no differences in percent of time spent in each of the four regions (outer, outer-inner; center-outer and center; Fig. 4c).

**Figure 4.**
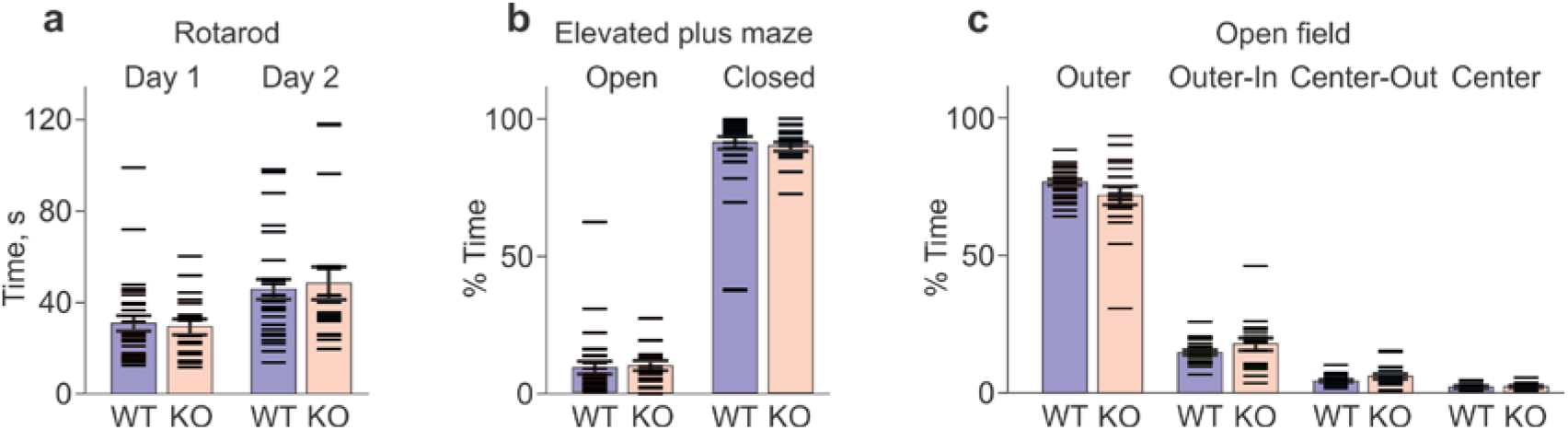
A1R-KO mice show no impairment in motor learning, nor increased anxiety compared to WT controls. In **a-c**, data from each animal (horizontal dash symbols), and group averages with SEM are shown. **a**: Latency to fall from an accelerating rotation drum (rotarod test) in WT and KO mice on days 1 and 2. Latency increased on day 2 in both, WT (from 31.1±3.4s to 45.9±4.4s, N=30, p<0.001) and KO groups (from 29.5±3.3s to 48.7±7.2s, N=18, p=0.0011). No difference between WT and KO (p=0.76 for day 1; p=0.74 for day 2). **b**: Percent time spent on the open and on the closed arm of the elevated plus maze. No difference between WT and KO groups (Open arm: 9.0±2.3% *vs.* 10.1±1.7%, p=0.71; Closed arm: 91.0±2.3% *vs.* 89.9±1.7%, p=0.71). **c**: Percent time spent during the open field test in the four virtually defined regions: Outer, Outer-In, Center-Out and Center. No difference WT *vs.* KO groups for any of the quadrants: Outer 78.1±1.06% *vs.* 73.2±3.44%, p=0.11; Outer-In 15.1±0.75% *vs.* 18.2±2.33%, p=0.13; Center-Out 4.8±0.33% *vs.* 6.3±0.97%, p=0.1; Center 2.0±0.18% *vs.* 2.3±0.31%, p=0.35.

Thus, KO animals showed no motor deficits or increased anxiety compared to WT mice. The absence of confound was further supported by analysis of a linear model of visual discrimination performance including additional tasks (response: Percent Correct days 14-19; predictors: Genotype, Sex, Age, Rotarod, Elevated plus maze, and Open field scores; F_DF(9,38)_= 3.477, p=0.0032). The only significant predictor of performance on the pairwise discrimination task remained Genotype (t = 3.366, p=0.0018; for any other predictor p>0.1), and predictor subsets optimized to minimize residual standard error always included Genotype.

Collectively, results showed that deletion of A1Rs selectively impaired repetitive learning on consequent visual tasks, but not learning on initial visual tasks, nor overall motor function or anxiety level of KO subjects.

### Lack of A1Rs in visual cortex of knockout mice

Finally, we verified that KO mice indeed lack A1Rs in visual cortex neurons. It is well established, that activation of A1Rs with 20 μM adenosine reliably suppresses synaptic transmission in visual cortex (Bannon et. al., 2014; Zhang et. al., 2015; van Aerde et. al., 2015; Yang et. al., 2020). We tested effects of adenosine using slices from plasticity experiments, and from a subset of behaviorally tested animals. In all tested WT neurons, 20 μM adenosine suppressed EPSPs (Fig. 5a, 46.7% of control; Fig. 5b, mean 44.9±2.6%, N=9). In contrast, 20 μM adenosine did not suppress EPSP amplitudes in any tested KO neuron (Fig. 5c, 97.4% of control; Fig. 5b, mean 98.6+1.3%, N=27). These results provide physiological verification of genotyping and confirm the absence of A1Rs in visual cortex of KO mice.

**Figure 5.**
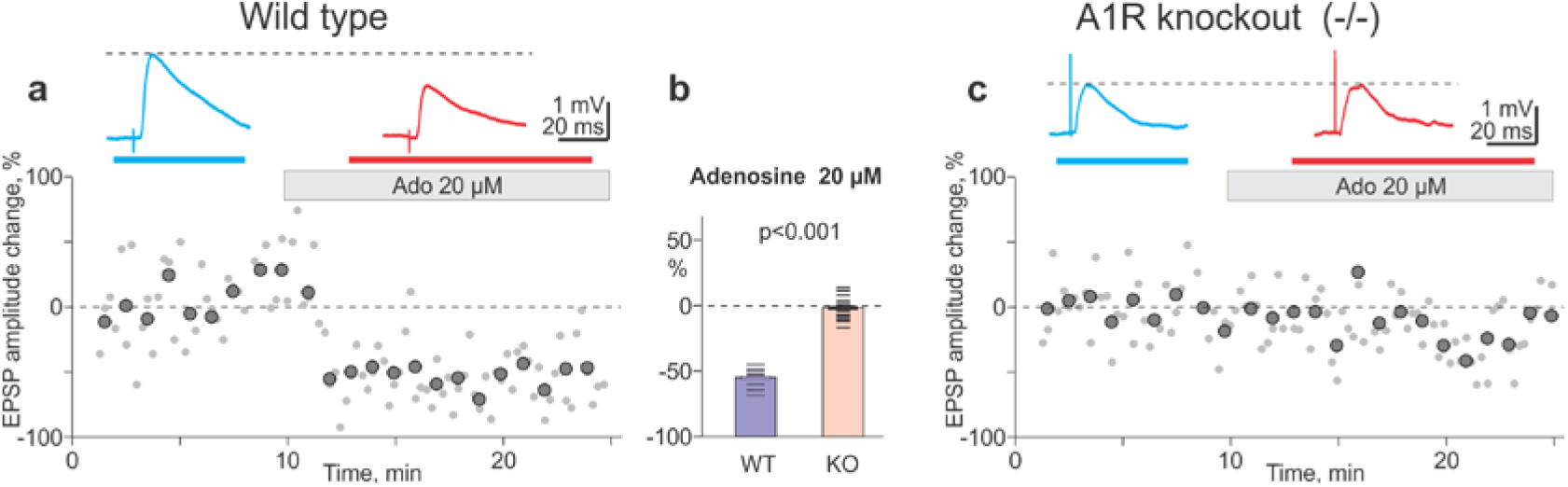
Physiological verification of A1R deletion in knockout mice: Adenosine suppresses synaptic transmission in visual cortex neurons from WT, but has no effect in A1R-KO mice. **a, c**: Time course shows individual EPSP amplitudes (grey dots) and averages over 1 min (large symbols) in two example neurons from WT (**a**) and A1R-KO (**c**) animals before and during application of 20 μM adenosine (grey horizontal bars). **b**: Changes of EPSP amplitudes in adenosine; data for individual neurons (horizontal dash symbols) and averages for WT (N=9 cells) and KO (N=27 cells) animals.

## DISCUSSION

Our results show impaired synaptic plasticity in visual cortex neurons, and deficits in visual learning, in A1R-KO mice compared to WT animals. Observed deficits were not. ‘all-or-none,’ but subtle and task-specific. Synaptic plasticity – both homosynaptic and heterosynaptic – could still be induced in visual cortex neurons from KO animals. However, there was a clear shift in the balance between LTP and LTD toward depression. Behaviorally, A1R-KO mice could still learn, and on a simple visual task they learned as well as WT subjects. However, on subsequent tasks of increasing difficulty, visual learning deficits in KO animals became progressively apparent. The most difficult test of pairwise visual discrimination revealed a dramatic impairment of learning in KO animals compared to WT controls.

### Experimental results support predictions from computer simulations

This observed dissociation between intact initial learning, and severe deficits in subsequent learning (or re-learning), was predicted to follow from compromised heterosynaptic plasticity (Volgushev et. al., 2016). Our simulations showed that model neurons and networks equipped with Hebbian-type learning rules and experimentally-observed heterosynaptic plasticity could learn to discriminate input patterns, and then repeatedly re-learn to discriminate new patterns. In contrast, models equipped with only Hebbian-type but no heterosynaptic plasticity could learn the first pattern discrimination, but re-learning was impaired. With each new subsequent task, the impairment became more severe due to runaway dynamics and eventual saturation of synaptic weights (Volgushev et. al., 2016) – a known drawback of Hebbian-type learning rules (Oja 1982; MIller and MacKay 1994; van Ooyen 2001; Zenke et. al., 2013). In a follow-up study we found that adenosine, acting via A1Rs, modulates heterosynaptic plasticity (Bannon et. al., 2017). In model neurons, heterosynaptic plasticity associated with functional A1Rs supported a homeostatic regime, bringing excessively changed synaptic weights back into the operating range (Bannon et. al., 2017). We predicted that this would “prepare” neurons for new learning. In contrast, impairment of heterosynaptic plasticity via blockade of A1Rs disrupted the homeostatic regime, and we predicted this would subvert capacity for re-learning. Results from the present study conform precisely to our predictions. At the synaptic level, lack of A1Rs in KO neurons impaired heterosynaptic plasticity and homeostatic regime, hindering the preparation of neurons for the next round of learning. At the behavioral level, this was associated with an impaired ability for progressive learning on new behavioral tasks in KO mice.

The need for heterosynaptic plasticity in learning systems equipped with Hebbian-type learning rules has been long appreciated in theoretical and modeling studies (Oja 1982; MIller and MacKay 1994; van Ooyen 2001; von der Malsburg 1973; Miller 1996). Moreover, theoretical work demonstrated that details of the mechanisms of heterosynaptic plasticity can influence learning, e.g. specifics of synaptic weight normalization determine the ability of a system to learn discrimination of subtle differences in input patterns (Oja 1982; MIller and MacKay 1994; Miller 1996). However, the role of heterosynaptic plasticity in learning has escaped experimental analysis, largely due to a lack of tools for selective manipulation. Here, we circumvent this barrier by using our prior experimental and theoretical analysis to generate specific predictions about behavioral consequences of experimentally observed modulation of heterosynaptic plasticity. Testing these predictions allowed us, to the best of our knowledge for the first time, to link an impairment of heterosynaptic plasticity to a behavioral deficit in re-learning on consecutive tasks. **This provides the first evidence for the role of heterosynaptic plasticity, and its postulated requirement for homeostatic synaptic function, in organism-level learning.**

### Regional specificity of A1R-function

Earlier studies using A1R-KO mice reported no impairment of synaptic plasticity in the hippocampus and no deficits in spatial learning, including reversal and working memory tests (Giménez-Llort et. al., 2002). Here we report contrasting results for the visual system: both synaptic plasticity in visual cortex, and the ability to re-learn visual tasks, was impaired in A1R-KO mice. These contrasting results indicate that, despite established similarities between synaptic plasticity in visual cortex and hippocampus (Kirkwood et. al., 1993), details of plasticity modulation – in this case by A1Rs – might be brain region-specific. At the same time, both hippocampus and visual cortex show a correspondence between impaired synaptic plasticity (or absence of impairment) and the ability to learn on respective tasks.

### Exclusion of confounds in behavioral learning results

Initial studies also reported that A1R-KO mice have decreased muscle strength, but no impairment in motor coordination, and increased anxiety compared to WT animals (Giménez-Llort et. al., 2002; 2005; Johansson et. al 2001). With the use of a different (rotarod) test, we confirmed normal motor learning and coordination in KO animals. However, we did not find increased anxiety in KO animals. The discrepancy could be due to the use of different tests (dark-light box and elevated plus maze with transparent walls in (Giménez-Llort et. al., 2002; Johansson et. al 2001), *vs.* open field and elevated plus maze with non-transparent walls in our study), and requires further testing.

Several lines of evidence indicate that impairment of learning on visual discrimination task in A1R-KO mice was not due to general functional deficits, such as poor vision, motor function or altered levels of anxiety or motivation. KO mice can see, because they learned simple visual tasks at a level equivalent to WT mice. Motor deficits could not explain the observed impairment of learning in A1R-KO animals because (i) the motor component of all three visual tasks was the same; (ii) performance on rotarod and open field tests was comparable in KO and WT mice, and did not predict learning outcomes; and (iii) the total number of responses (correct and incorrect) during learning on the third visual task was the same in WT and KO mice. The same total number of responses argues against differences in physical fatigue or impaired motivation in WT and KO subjects. Such impairments would typically manifest in reduced responses and/or trials completed. Comparable number of responses also argues against an increased level of anxiety in KO mice, together with evidence of comparable performance on an elevated plus maze, and failure of plus-maze results to predict learning outcomes. Overall, we conclude that observed deficits in learning visual tasks in A1R-KO animals were not due to general functional deficits, but reflect specific impairment of synaptic plasticity in visual cortex neurons.

### Conclusions and Outlook

The present study provides, to our knowledge, the first experimental evidence for a link between impaired heterosynaptic plasticity and a specific behavioral deficit – progressive impairment of learning on consecutive tasks. We previously predicted that changes in heterosynaptic plasticity following A1R blockade would lead to such a specific learning deficit (Bannon et. al., 2017; Volgusuev et. al., 2016). Experimental results confirming this prediction offer broader evidence in support of the proposed homeostatic role of heterosynaptic plasticity during on-going associative learning (Oja 1982; MIller and MacKay 1994; von der Malsburg 1973; Miller 1996; Watt et. al., 2010; Chistiakova et. al., 2015; Zenke and Gerstner 2017; Bannon et. al., 2020). Our novel experimental evidence for the role of heterosynaptic plasticity in learning opens up a whole new range of questions. From an experimental perspective, our data invite the use of specific tools for manipulating A1R-mediated modulation of heterosynaptic plasticity (e.g., conditional, region-specific or cell-type specific knockout models, or local and time-restricted A1R-blockade) to interrogate constraints on the requirement for heterosynaptic plasticity for repetitive learning. Another important question is specificity of the A1R-mediated modulation of homeostatic function of heterosynaptic plasticity with respect to brain region and sensory modality subserving learning (e.g., auditory or tactile learning). Also, research using brain regions and learning tasks for which A1R-deletion is not critical (e.g., homosynaptic plasticity and spatial learning in the hippocampus (Giménez-Llort et. al., 2002; 2005)), might reveal further mechanisms that regulate synaptic homeostasis during associative learning. The link between heterosynaptic plasticity and the ability for repetitive learning also provides opportunity to examine a putative role for A1R-modulation of heterosynaptic plasticity in state-dependence of learning across sleep-wake cycles (Tononi and Cirelli 2006; 2014; Bannon et. al., 2017).

A final intriguing question concerns whether heterosynaptic plasticity could be selectively upregulated *in vivo* to support the homeostatic regime. Such targeted interventions could profoundly alter and enhance learning. They could also lead to therapies for brain disorders associated with excessive potentiation of pathologic connectivity (e.g., epilepsy, PTSD and chronic pain). Such interventions could capitalize on established modulation of plasticity via adenosine/A1R (Bannon et. al., 2017, and present results), but could also be expanded to other synaptic modulators, offering new therapeutic avenues.

## ACKNOWLEDGEMENTS

The work on this project was supported by a grant from the University of Connecticut Research Excellence Program to MV and RHF, support from IBACS to the Murine Behavioral Neurogenetics Facility and RHF, and a grant from the Russian Science Foundation #20-15-00398 to AM. We are grateful to Margaret Balogh, Emily Gombotz and Gabriel Vega for help with some of the experiments.

## AUTHOR CONTRIBUTIONS

Designed research: MV, RHF, AM; conducted experiments: RC, AM, MV; processed data, prepared figures and drafted the manuscript RC, AM, RHF, MV.

## METHODS

All experimental procedures in this study were re in compliance with the US National Institutes of Health regulations and were approved by the Institutional Animal Care and Use Committee of the University of Connecticut.

### Subjects

We used A1R knockout mouse strain B6N.129P2-*Adora1*^tm1Bbf^/J obtained from The Jackson Laboratory (stock No 014161, Cryo Recovered; https://www.jax.org/strain/014161) to establish breeding colony at the University of Connecticut animal facilities. Genotyping was made by Transnetyx (Cordova, TN, USA; https://www.transnetyx.com/). For experiments, we used A1R (−/−) knockout (A1R KO) and littermate wild type (WT) animals of both sexes.

### Electrophysiology

#### Preparation of slices

Details of slice preparation and recording are similar to those used in previous studies (Lee et. al., 2012; Volgushev et. al., 2016; Bannon et. al., 2017), but using sucrose-based solution during slice preparation. Adult mice (78 – 188 days old, both sexes, WT or A1R KO) were anaesthetized with isoflurane, decapitated, and the brain quickly removed and placed into an ice-cold oxygenated solution containing, in mM: 83 NaCl, 25 NaHCO_3_, 2.7 KCl, 1 NaH_2_PO_4_, 0.5 CaCl_2_, 3.3 MgCl_2_, 20 glucose, 71 sucrose, bubbled with 95% O_2_/5% CO_2_. Coronal slices (350 μm thickness) containing the visual cortex were prepared from the right hemisphere and placed in a slice incubator filled with oxygenated sucrose-based solution. After slices recovered for 45-60 min at 34°C, slice incubation chamber was moved to room temperature. For recording, individual slices were transferred to a recording chamber mounted on an Olympus BX-50WI microscope equipped with IR-DIC optics. Recordings were made in a solution containing, in mM: 125 NaCl, 25 NaHCO_3_, 25 glucose, 3 KCl, 1.25 NaH_2_PO_4_, 2 CaCl_2_, 1 MgCl_2_, bubbled with 95% O_2_/5% CO_2_, pH 7.4, at 30°-32°C.

#### Intracellular recording and synaptic stimulation

Whole-cell recordings were made from Layer 2/3 pyramidal cells from visual cortex. Intracellular pipette solution contained, in mM: 130 K-Gluconate, 20 KCl, 10 HEPES, 10 Na-Phosphocreatine, 4 Mg-ATP, 0.3 Na2-GTP, (pH 7.4 with KOH). Monosynaptic excitatory postsynaptic potentials (EPSPs) were evoked using two pairs of bipolar stimulating electrodes (S1 and S2) placed in layer 4, below the L2/3 recording site. Stimuli were applied to S1 and S2 in alternating sequence, so that each input was stimulated each 15 seconds. To test for the possible contribution of inhibition, evoked PSPs were recorded at depolarized potentials between −50 and −40 mV. Only those PSPs that were still depolarizing at this membrane potential were considered excitatory and included in the analysis. Membrane potential and input resistance were monitored throughout experiments; cells in which either parameter changed by more than 15% by the end of recording were discarded.

#### Plasticity induction

Synaptic plasticity was induced by either a pairing procedure (STDP protocol) or intracellular tetanization. During the pairing procedure, EPSP evoked at one of the two independent inputs was followed with a 10 ms delay by a burst of five spikes evoked by depolarizing pulses (5 ms, 100 Hz; see Fig. 1 inset). Pairing was repeated 30 times, in three trains (1/min) of ten episodes (1 Hz). Intracellular tetanization consisted of the same pattern of postsynaptic activation: three trains (1/min) of ten bursts (1 Hz) of five action potentials evoked by depolarizing pulses (5 ms, 100 Hz), but without synaptic stimulation (see Fig. 2A).

#### Behavioral testing

Behavioral testing on operant learning of visual tasks of increasing complexity was performed using the automated Bussey-Saksida touchscreen chambers (Campden Instruments Ltd, Loughborough, UK). Motor function and anxiety were assessed using Rotarod, Elevated plus maze and Open field tests.

During behavioral testing, all subjects were single-housed in standard mouse tubs under a 12h/12h light/dark cycle, food and water ad libitum. Two weeks before the start of operant training on visual task subjects were gradually transitioned to a restriction of 85% from their baseline weight. During the last week before training, subjects were given a sample (~1 ml) of the liquid food reward (Strawberry Ensure Plus, Abbott Nutrition, Columbus, OH) in their home cage. After completion of testing on visual learning task, animals were returned to ad libitum food and water. All behavioral testing occurred during the light cycle and performed blind to genotype.

#### Visual learning task

All training and testing sessions were performed using the automated Bussey-Saksida touchscreen chambers (Campden Instruments Ltd, Loughborough, UK) which had a trapezoidal operant area, a touchscreen (30.7 cm, resolution 800 × 600), and a feeder situated across from the center of the screen. Visual stimuli were high contrast, large (size about 10% of the screen) clearly distinct geometric figures, presented in a pseudo-random order in the lower-right or lower-left quadrant of the screen. Each subject had one training session (60 min or until a maximum of 30 rewards is reached) per day. Operant learning consisted of pre-training, followed by three stages of learning visual tasks of increasing difficulty. During pre-training (two weeks), the mice learned to associate screen presses with reward delivery.

During the Stage 1 (‘must initiate’, 5 days), the subjects learned to initiate presentation of a visual stimulus on the screen by nose-poke and exit the reward tray, and then to touch the stimulus to obtain food reward. At this stage, touching other-than-stimulus part of the screen did not cause any actions. The number of obtained food rewards (‘correct responses’), as well as total duration of the session (time it took to obtain 30 rewards, or 60 min), were recorded.

During Stage 2 (‘punish incorrect’, 5 days) the subjects learn to touch only the stimulus and not any other part of the screen. Touching the stimulus (‘correct’) is rewarded with food. Touching any other, blank, portion of the screen (‘incorrect’) is punished by a time out for 5 seconds, during which no inputs are registered and the test cage is illuminated with bright light (~60 lx). Number of correct responses, number of incorrect responses and duration of the session were recorded.

During Stage 3 (‘pairwise discrimination’, 19 days), two visual stimuli are presented on the screen, and the subjects learn to press a correct stimulus. Touching correct stimulus is rewarded with food. Touching incorrect stimulus or blank part of the screen is punished as above, by a 5 seconds time out and the test cage illuminated with bright light (~60 lx). Number of correct responses, number of incorrect responses and duration of the session were recorded.

#### Motor function and learning: Rotarod test

Subjects were placed on a rotating drum that gradually accelerated from 4 to 40 rotations per minute across a span of 2 minutes. Latency for mice to fall from the rotating drum was recorded. Subjects were tested for two consecutive days, four tests per day.

#### Motor activity and anxiety: Elevated plus maze and open field tests

In the Elevated Plus Maze test subjects were placed in the middle of an elevated cross with two arms opposite to each other having two high side walls (“closed arm”) and the other two arms having no walls (“open arm”). Mouse movement was monitored over five minutes using TopScanLite (CleverSys, Reston, VA), and time spent in the open and in the closed arm, as well as the number of entries into each arm were recorded.

In the Open Filed test subjects were placed in the center of a square box with high side walls and no top (50 cm × 50 cm × 50 cm), and their movement was monitored for 15 minutes. Time spent in each of the four virtually defied regions: outer, outer-inner, center-outer and center was recorded using TopScanLite (CleverSys, Reston, VA).

#### Data processing and statistical analyses

*Electrophysiological data* analysis was made using custom-written programs in MatLab (© The MathWorks, Natick MA, USA), Excel (MS Office 2010) and scripts in R (The R Foundation for Statistical Computing version 3.4.0, 2017-04-21). All inputs included in the analysis fulfilled the following criteria (1) excitatory nature of EPSP, as verified by the absence of reversal when recorded at depolarized potentials between −40 and −50 mV; (2) stability of EPSP amplitudes during the control period, (3) stability of the membrane potential and input resistance throughout the recording, and (4) stability of the onset latency and kinetics of the rising slope of the EPSP. Amplitudes of EPSPs were measured as the difference between the mean membrane potential during two time windows, the first time window placed before the onset and the second window placed on the rising slope of the synaptic response, just before the peak. For calculating significance of response amplitude changes at individual inputs, T-test was used to compare control responses recorded before the application of plasticity-induction protocol (n=15-35 responses from a stationary period just before plasticity induction) to responses after plasticity induction (n=40-120 responses from a period typically around 20-60 min after the induction). Response changes (LTP or LTD) were considered significant at p<0.05. For calculation of population averages across inputs, response amplitudes in each input were first normalized to control, and then averaged across inputs. For comparison of frequency of occurrence of LTP and LTD Chi-square test was used.

*Behavioral data* analysis was made using Excel (MS Office 2010) and scripts in R (The R Foundation for Statistical Computing, version 3.4.0, 2017-04-21). Only subjects tested on all behavioral tests described above were included in the final analysis. We have excluded one subject (KO, male) who initiated only few trials on any day during training on visual task (<10 correct and incorrect trials on most of the days and never reached 20; other subjects completed 38.2±0.4 trials per day, gross average over all days and subjects). Behavioral results presented in this study were obtained from N=18 KO (13F, 5M) and N=30 WT (11F, 19M) subjects. We used t-test for comparison of values and Chi-Square test for comparison of frequency of occurrence in pairs of samples. For analysis of interaction between multiple variables we used linear model analysis employing functions *regsubsets* and *lm*, and scripts in R.

Throughout the text, averages are given with SEM; p-values >0.001 are given in full and p-values <0.001 as p<0.001.

### Resource Availability

#### Lead Contact

Further information and requests for resources and reagents should be directed to and will be fulfilled by the Lead Contact, Maxim Volgushev.

#### Materials Availability

This study did not generate new unique reagents.

#### Data and Code Availability

Data for summary figures in the paper are provided in the supplementary tables. Original data and processing codes are available from the corresponding author on request.

---

**SUPPLEMENTAL INFORMATION: DATA FOR FIGURE 1**

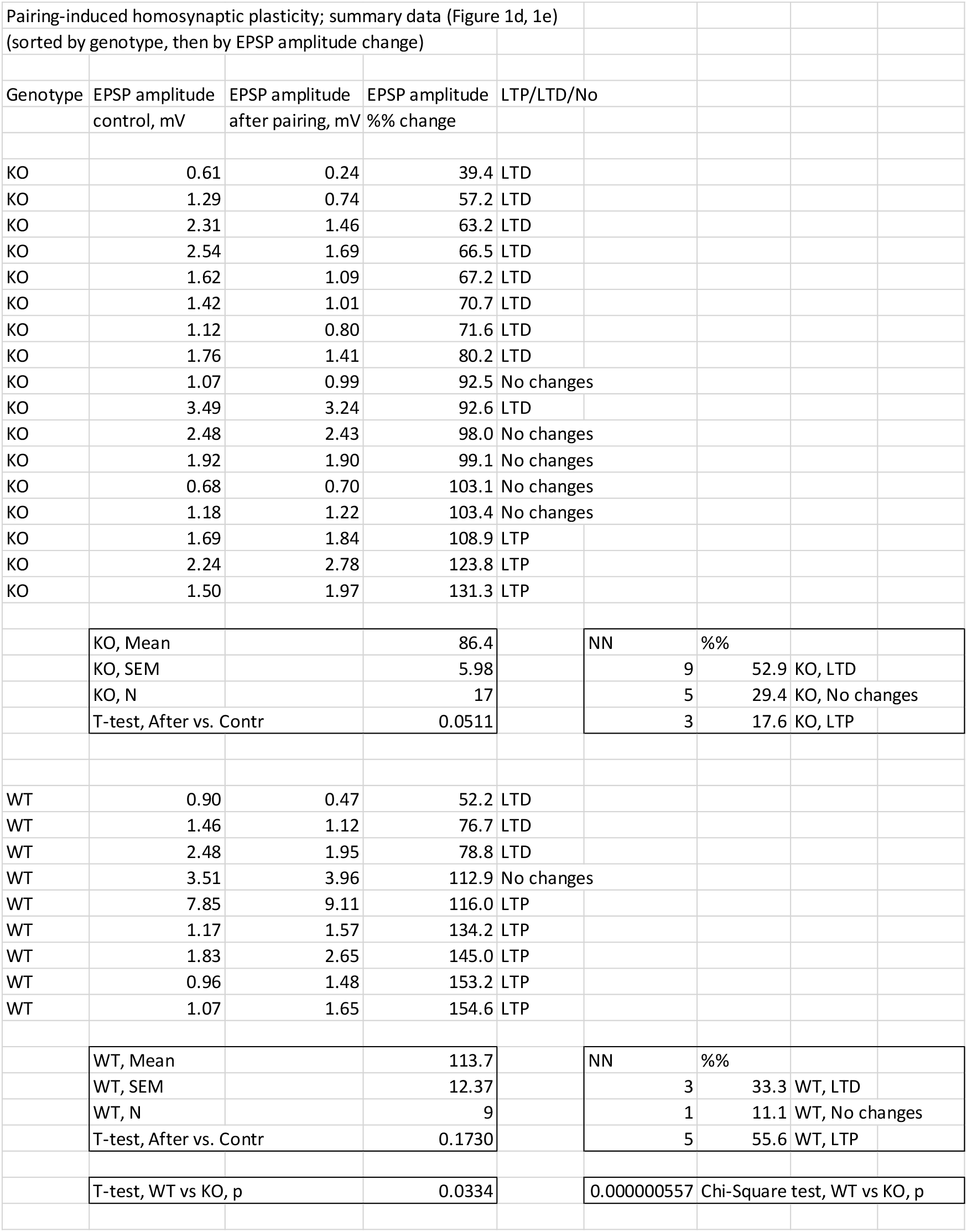

**SUPPLEMENTAL INFORMATION: DATA FOR FIGURE 2**

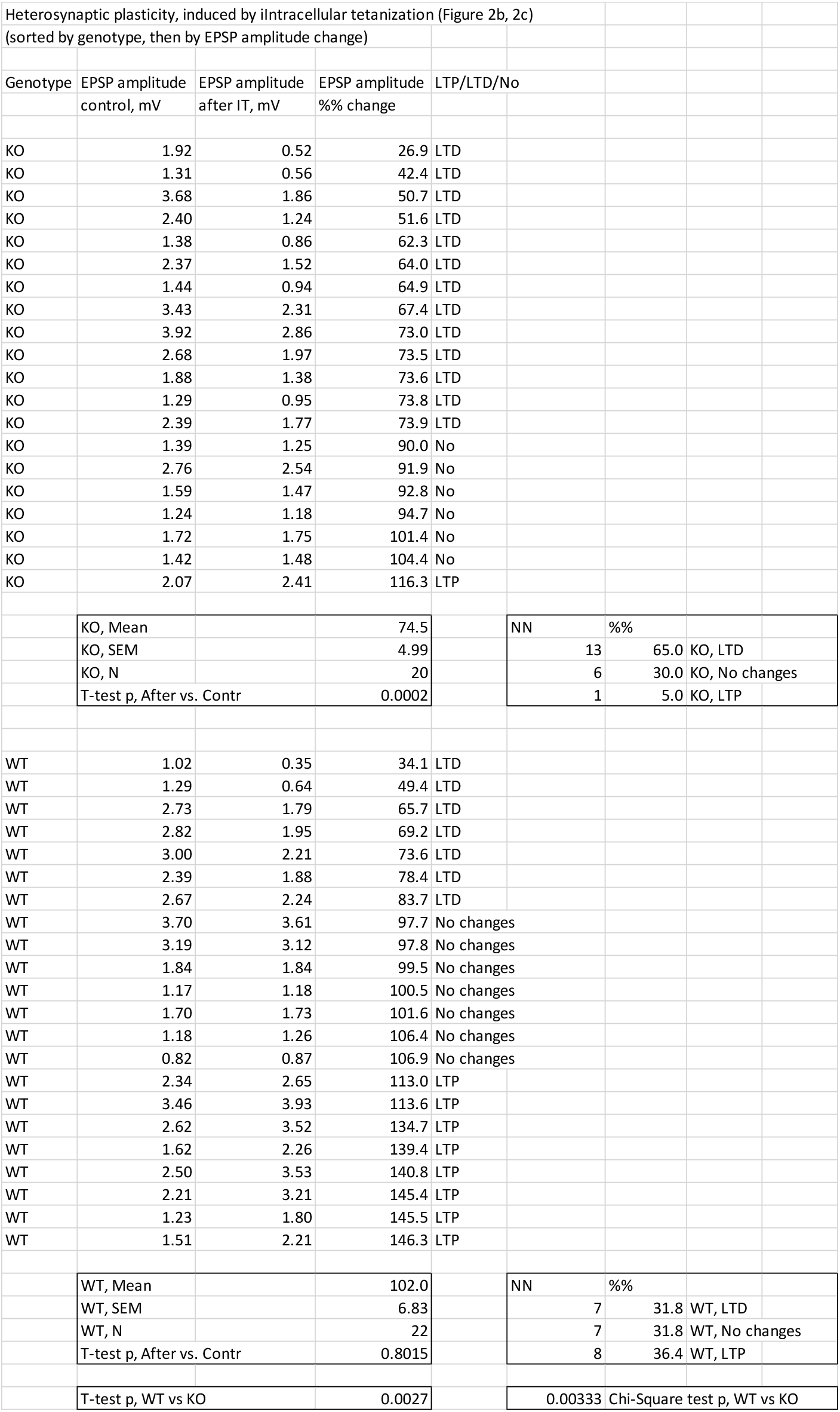

**SUPPLEMENTAL INFORMATION: DATA FOR FIGURE 3 (part 1 of 4)**

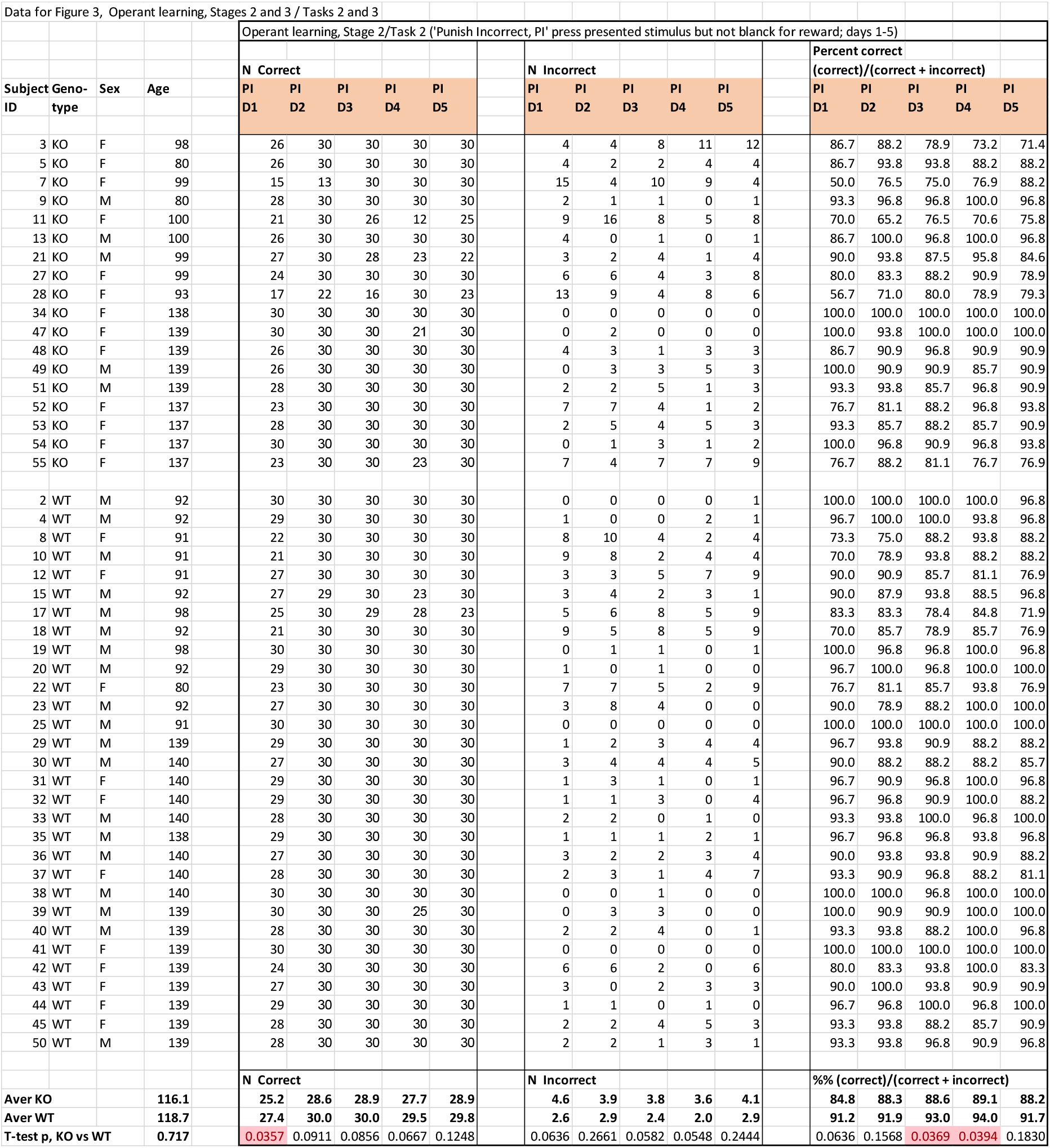

**SUPPLEMENTAL INFORMATION: DATA FOR FIGURE 3 (part 2 of 4)**

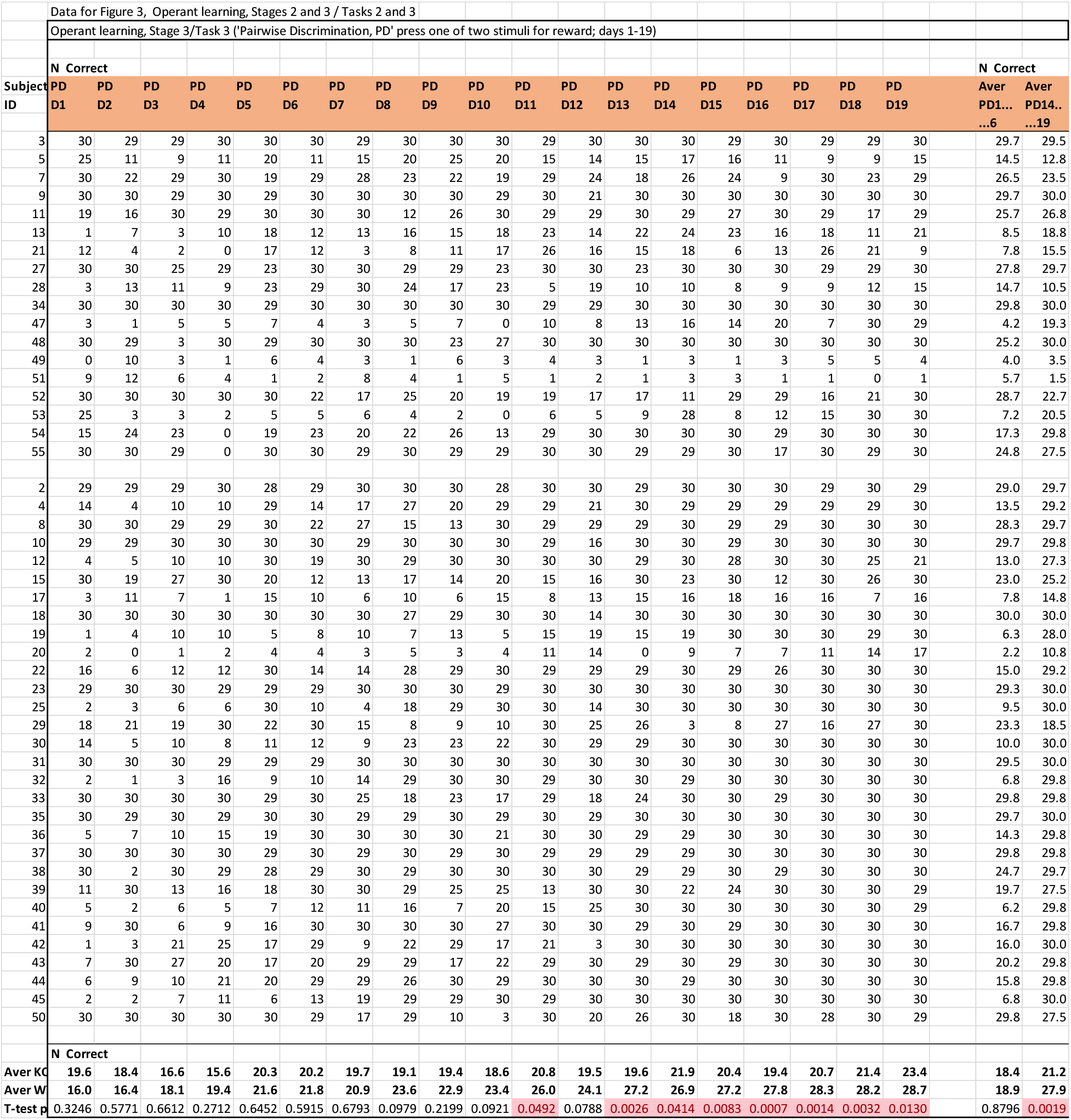

**SUPPLEMENTAL INFORMATION: DATA FOR FIGURE 3 (part 3 of 4)**

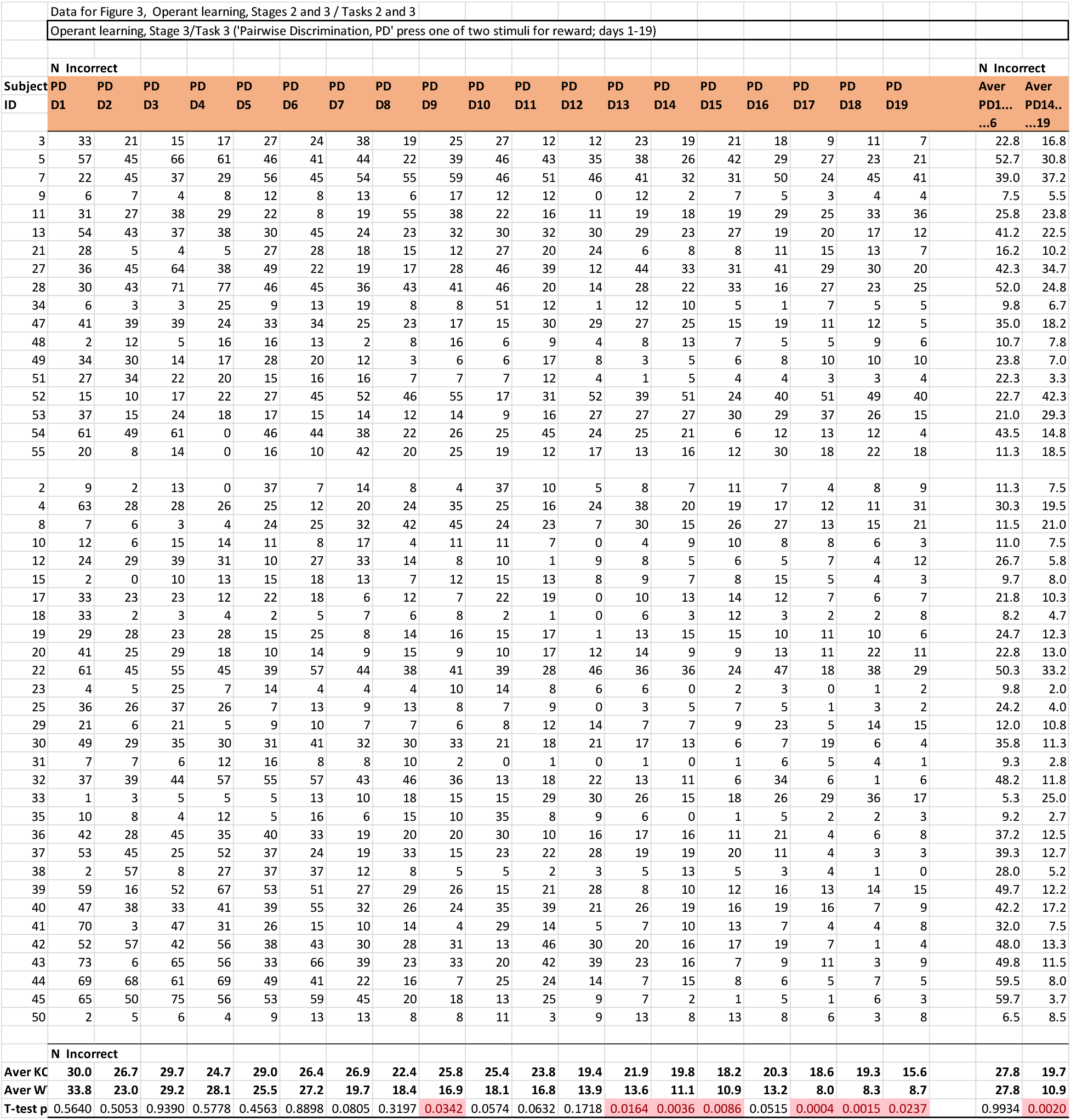

**SUPPLEMENTAL INFORMATION: DATA FOR FIGURE 3 (part 4 of 4)**

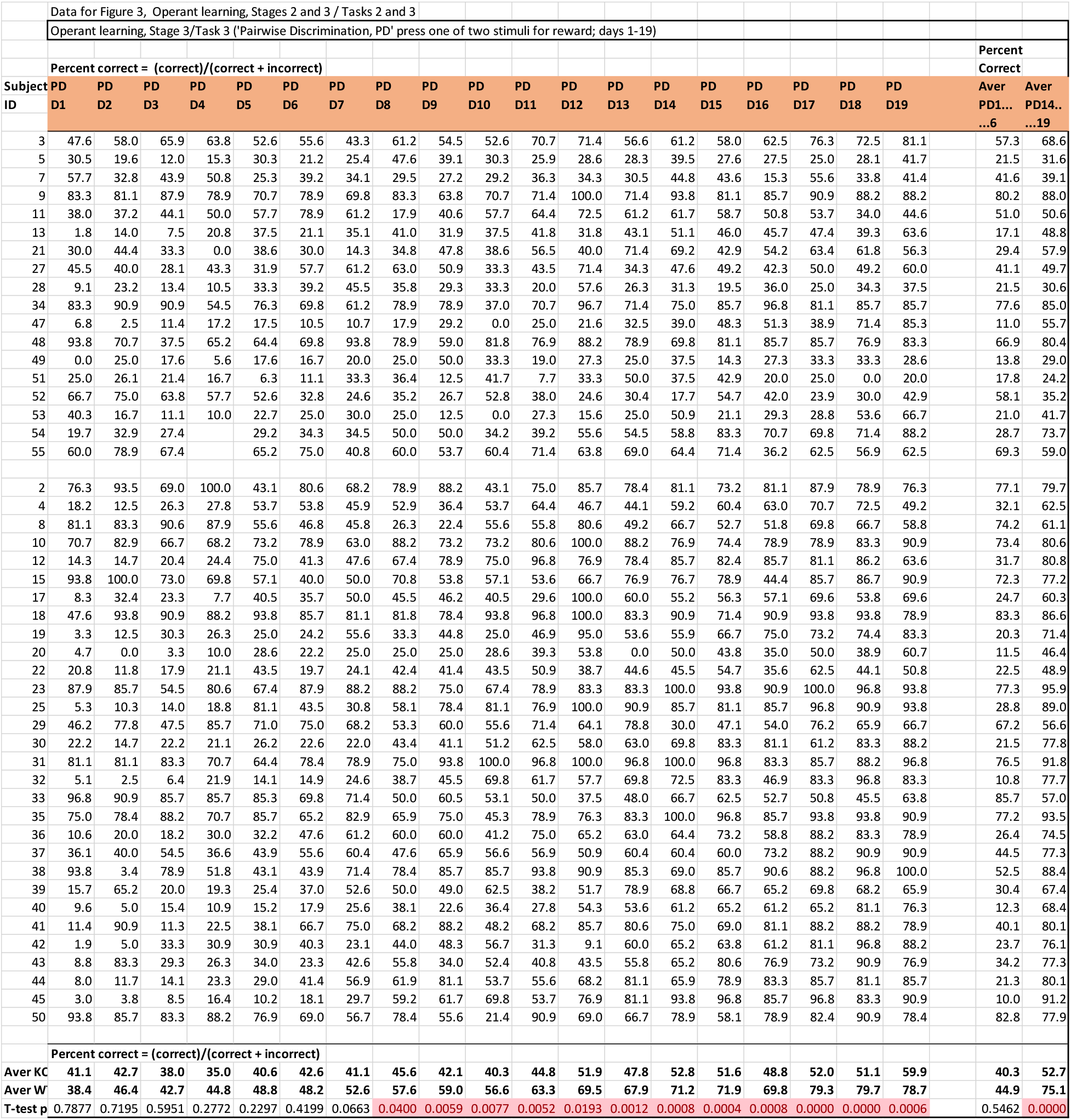

**SUPPLEMENTAL INFORMATION: DATA FOR FIGURE 4**

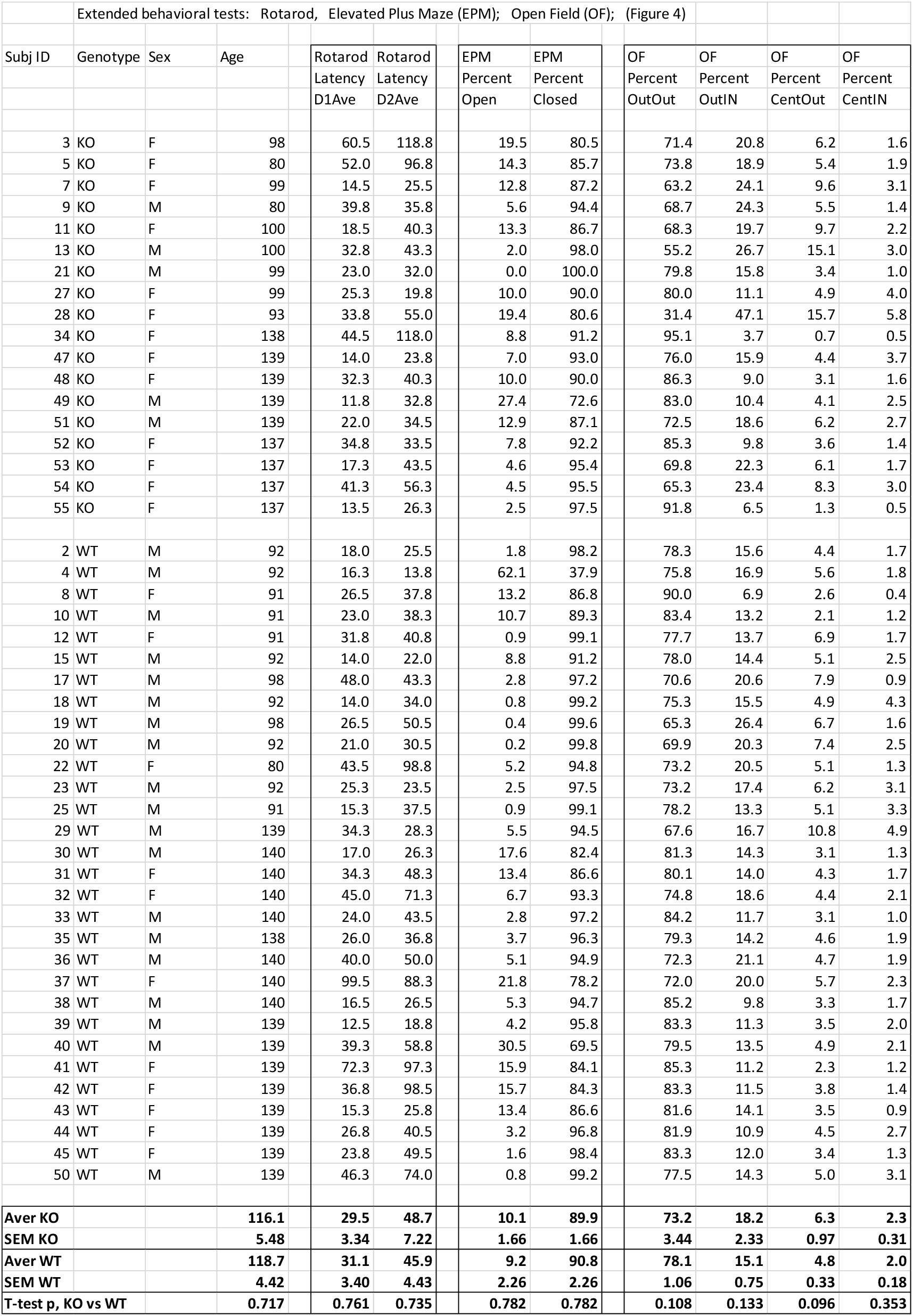

**SUPPLEMENTAL INFORMATION: DATA FOR FIGURE 5**

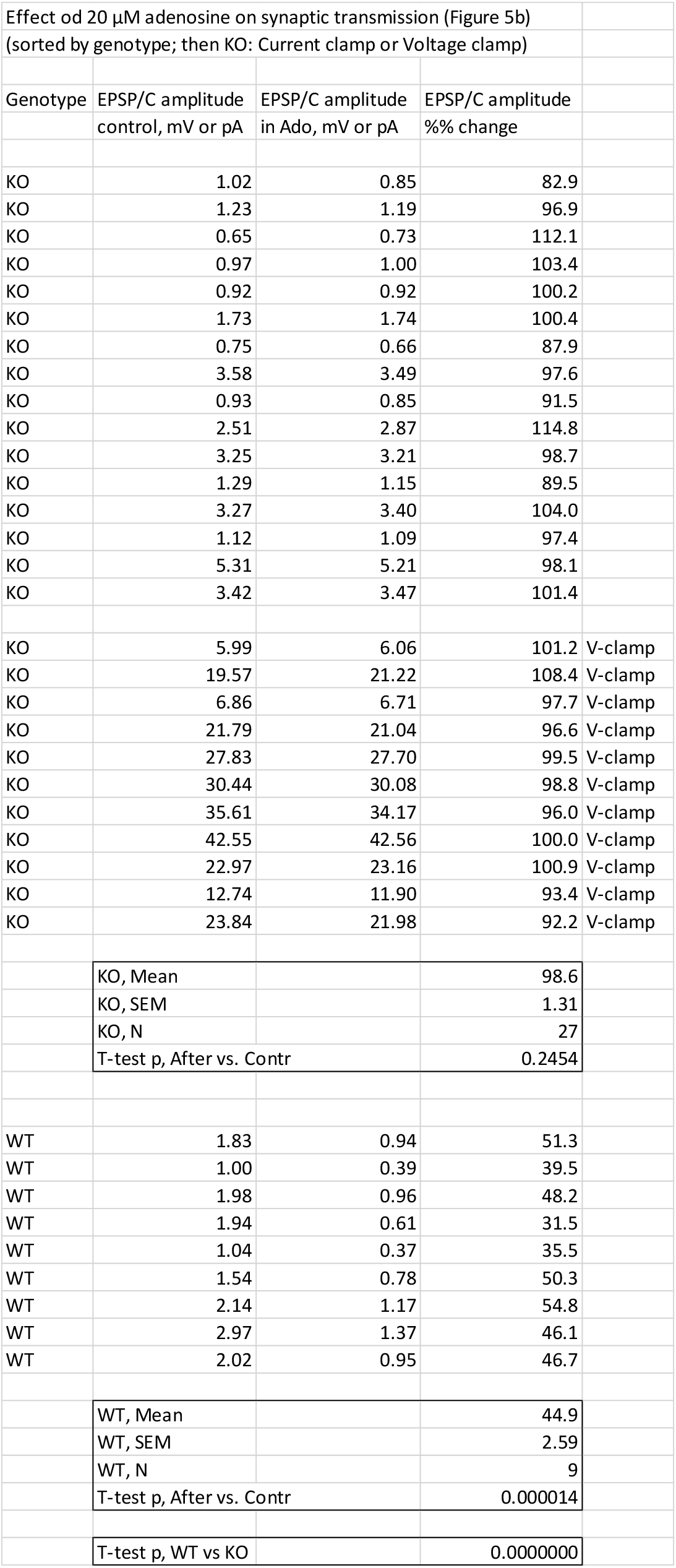

